# Sulcal depth in medial ventral temporal cortex predicts the location of a place-selective region in macaques, children, and adults

**DOI:** 10.1101/2020.01.27.921346

**Authors:** Vaidehi S. Natu, Michael J. Arcaro, Michael A. Barnett, Jesse Gomez, Margaret Livingstone, Kalanit Grill-Spector, Kevin S. Weiner

**Author notes:** **Corresponding Author:** Vaidehi S. Natu, Department of Psychology, Stanford University, Stanford.

## Abstract

The evolution and development of anatomical-functional relationships in the cerebral cortex is of major interest in neuroscience. Here, we leveraged the fact that a functional region selective for visual scenes is located within a sulcus in medial ventral temporal cortex (VTC) in both humans and macaques to examine the relationship between sulcal depth and place-selectivity in medial VTC across species and age groups. To do so, we acquired anatomical and functional magnetic resonance imaging scans in 9 macaques, 26 human children, and 28 human adults. Our results revealed a strong structural-functional coupling between sulcal depth and place-selectivity across age groups and species in which selectivity was strongest at the deepest sulcal point (the sulcal pit). Interestingly, this coupling between sulcal depth and place-selectivity strengthens from childhood to adulthood in humans. Morphological analyses suggest that the stabilization of sulcal-functional coupling in adulthood may be due to sulcal deepening and areal expansion with age as well as developmental differences in cortical curvature at the pial, but not the white matter surfaces. Our results implicate sulcal features as functional landmarks in high-level visual cortex and highlight that sulcal-functional relationships in medial VTC are preserved between macaques and humans despite differences in cortical folding.

## Introduction

Primate visual cortex comprises a few dozen visual areas spanning occipital, temporal, and parietal cortices (Felleman and Van Essen 1991; Van Essen et al. 2001; Rosa and Tweedale, 2005). Although the topological layout of these areas is similar across individuals within a particular species (Tsao et al. 2003; Grill-Spector and Weiner 2014; Arcaro and Kastner 2015; Arcaro and Livingstone 2017; Larsson and Heeger 2006; Dumoulin et al. 2000; Witthoft et al. 2014; Kolster et al. 2014), the shape and size of brains varies considerably across species (Sereno and Tootell 2005; Zilles et al. 2013). Despite this variability, there is a strong correspondence between sulcal folding and the functional organization of primary visual cortex (V1). In both humans and monkeys, V1 is localized to the calcarine sulcus (Tootell et al. 1998; Hinds et al. 2008; Rajimehr and Tootell 2009; Benson et al. 2012) and surface topology is predictive of the underlying retinotopic representations (Rajimehr and Tootell 2009; Benson et al. 2012). Beyond V1, the traditional view is that there is little correspondence between sulcal folding and visual areas given that (1) there are more brain areas than there are sulci, (2) the number of sulci varies across primates, and (3) there is a relatively large degree of individual variability in the size and topological layout of cortex beyond primary sensory areas (Van Essen et al. 2019; Zilles et al. 2013).

Nevertheless, the last decade has shown renewed interest in the relationship between sulcal folding and functional regions (Amiez et al. 2019; Amiez et al. 2018; Amiez and Petrides 2014, Weiner et al. 2014; Witthoft et al. 2014; Benson et al. 2012, 2014; Leroy et al. 2015; Bodin et al. 2018; Lopez-Persem et al. 2019; Weiner 2019). There is a surprising level of orderliness between sulcal folds and functional representations in late-maturing ventral temporal cortex (VTC) of humans (children and adults; Weiner et al. 2010; Weiner et al. 2011; Grill-Spector and Weiner 2014; Weiner et al. 2018; Natu et al. 2019) as well as macaques (Nasr et al. 2011; Arcaro and Livingstone 2017). Focusing on sulcal landmarks as predictors of human high-level visual regions has revealed a coupling between sulcal features (as opposed to entire sulci) and functional regions. For example, in human VTC, there is a strong predictable relationship between a) the anterior tip of the mid-fusiform sulcus and the location of a functional region selective to faces (Weiner et al. 2014) and b) a sulcal intersection of the collateral sulcus and the location of a region selective for visual places and scenes (Weiner et al. 2018).

Beyond large-scale sulcal folding patterns, there is also interest in morphological features of sulci such as the deepest point in a sulcus, which is known as the sulcal pit. The sulcal pit is interesting because it is hypothesized to be associated with functional specialization (Smart and McSherry 1986; Rakic 1988; Welker 1990; Regis et al. 2005; Hasnain et al. 2006; Lohmann et al. 2008; Im et al. 2010; Im et al. 2011; Meng et al. 2014; Auzius et al. 2015; Leroy et al. 2015; Bodin et al. 2018; Leroy et al. 2015; Im et al. 2019). In human adults, sulcal pits are shown to predict cognitive skills such as verbal IQ (Im et al. 2011) as well as the location of functional regions selective for human voices and social communication (Bodin et al. 2018). Critically, a stable spatial distribution of sulcal pits also exists at birth with regionally heterogeneous growth patterns in sulcal depth over the first 2 years of infant life (Meng et al. 2014; Meng et al. 2018). Thus, growing evidence suggests that morphology of sulcal folds, specifically sulcal pits, may have a tight coupling with functional regions – even in association cortices. Nevertheless, the relationship between sulcal depth and functional selectivity across development and species remains untested.

Here, we leveraged the anatomical-functional relationship between sulci in medial VTC and place-selectivity to examine the relationship between sulcal depth and functional selectivity across development and species. This place-selective region in medial VTC is interesting to study from a developmental perspective because it (i) emerges within 6 months postnatally in humans (Deen et al. 2017), (ii) it appears to be functionally mature in early childhood (Golarai et al. 2007; Scherf et al. 2007; Golarai et al. 2010; Gomez et al. 2017) while also exhibiting prolonged morphological development during childhood (Natu et al. 2019), and (iii) there is a proposed homologous place-selective region in medial VTC of macaques (Kornblith et al. 2013; Arcaro and Livingstone, 2017). To explore the relationship between place selectivity and sulcal morphology, we asked two main questions: (1) Is there a consistent anatomical-functional relationship between sulcal depth and the location of a place-selective region in human medial VTC? (2) Does this relationship change on developmental (childhood to adulthood) or evolutionary timescales? To answer these questions, we acquired anatomical and functional magnetic resonance imaging (fMRI) scans in 9 adult rhesus macaques (ages: 1-5 years, 4 female), 26 human children (ages 5-12, 17 female), and 28 human adults (ages 22-28, 14 female).

## Materials and Methods

### Participants

*Humans.* 26 children (ages 5-12, 17 female) and 28 adults (ages 22-28, 14 female) participated in our study. Children were recruited from the Palo Alto public school district through flyers and online advertisements. Adult participants are Stanford University affiliates. MRI data were collected using a 3T GE scanner in the Center for Cognitive and Neurobiological Imaging at Stanford University. All participants had normal or corrected-to-normal vision and provided written, informed consent. Protocols were approved by the Stanford Internal Review Board on Human Participants.

*Macaques.* 9 rhesus macaques (4 females, ages 1-5 years old) participated in our study. All procedures were approved by the Harvard Medical School Animal Care and Use Committee and conformed to National Institute of Health guidelines for the humane care and use of laboratory animals. Anatomical MRI data were collected using a 3T Siemens Skyra scanned in Brigham and Women’s hospital. Functional MRI data were collected using a 3T TimTrio scanner in the Athinoula A. Martinos Center for Biomedical Imaging of Massachusetts General Hospital.

### *Data acquisition* in humans (*Anatomical scans*)

For human subjects, participants were scanned using spin-echo inversion recovery with an echo-planar imaging (EPI), read-out (SEIR-EPI), with a 3T GE Signa scanner and a custom-built, phase array 32-channel, receive-only, head coil using methods as in prior studies (Mezer et al. 2013; Natu et al. 2016; Gomez et al. 2017; Gomez et al. 2018; Nordt et al. 2018; Natu et al. 2019). This scan was done with a slab-inversion pulse and spatial-spectral fat suppression. We used 2mm^2^ in-plane resolution with a slice thickness of 4 mm and the EPI readout was performed using 2X acceleration. QMRI parameters were measured from spoiled-gradient echo images acquired with different flip angles (α = 4°, 10°, 20° and 30°, TR = 14 ms, TE = 2.4 ms) and a voxel resolution of 0.8×0.8×1 mm, which was resampled to 1 mm^3^ isotropic. For SEIR-EPI, the TR was 3s. The echo time was set to minimum full; inversion times were 50, 400, 1200 and 2400 ms. Anatomical data were aligned to the AC-PC plane. From the qMRI data, we generated whole-brain T_1_-weighted anatomy. The spoiled-GE and the SEIR scans were processed using the mrQ software package in MATLAB to produce the T_1_ maps. The mrQ analysis pipeline corrects for RF coil bias using SEIR-EPI scans, producing accurate T_1_ fits across the brain. The full analysis pipeline can be found at (https://github.com/mezera/mrQ) (Mezer et al. 2013).

*Reconstruction of cortical surfaces.* Human T_1_ images underwent automated cortical surface reconstruction using FreeSurfer’s autosegmentation software (http://freesurfer.net). The anatomical images were segmented into white and gray matter. White matter surfaces were inspected and manually fixed for missing or mislabeled white matter voxels using ITK-SNAP (http://www.itksnap.org/). A mesh of each participant’s cortical surface was generated from the boundary of the white and gray matter and this mesh was inflated for visualization of functional activations inside the sulci.

*Functional scans were* obtained with the same scanner and coil using a T2*-sensitive gradient echo spiral pulse sequence with a resolution of 2.4 x 2.4 x 2.4 mm, TR = 1000 ms, TE = 30 ms, flip angle = 76°, and FOV = 192 mm. We collected 48 oblique slices, oriented parallel to the superior temporal sulcus, using a multiplexing technique allowing whole-brain coverage of functional data. During MRI scanning, participants were lying supine inside the magnet. Visual stimuli were projected onto a monitor and were viewed through an angled mirror mounted above the participant’s head. We defined place-selective regions of interest (ROIs) in individual participants using a functional localizer experiment based on our prior methods (Stigliani et al. 2015; Natu et al. 2016; Gomez et al. 2017; Gomez et al. 2018; Nordt et al. 2018; Weiner et al. 2018; Natu et al. 2019). Participants completed 3 runs of a functional localizer experiment (5.24 min/run) with 78, 4s blocks in each run. Participants viewed gray-scale stimuli, which were blocked by category. Images consisted of two subtypes from each of five categories: places (houses and indoor scenes), human faces (child faces and adult faces), characters (numbers and pseudo-words), bodies (limbs and headless bodies), and objects (guitars and cars). Each image was shown only once during the experiment. In each 4 s block, different stimuli of one of the above category were shown at a rate of 2 images per second. Blocks were counterbalanced across categories and also with baseline blocks consisting of a blank, gray background. *Oddball task:* During the scan, participants fixated on a central dot and pressed a button when a phase-scrambled oddball image appeared randomly in a block (∼33% of the blocks). Data of each human participant were corrected for within-run and between-run motion. Only runs with motion of less than two voxels were included in the study

*Analysis of functional data.* Localizer data were analyzed using code written in MATLAB with the mrVista toolbox (http://github.com/vistalab). Data were not spatially smoothed and were analyzed in each participant’s native brain space. The time courses of each voxel were converted from arbitrary scanner units into units of % signal change. For each participant, we ran a GLM to model each voxel’s time course. The experimental design matrix was convolved with the SPM hemodynamic response function (HRF) (http://www.fil.ion.ucl.ac.uk/spm) to generate predictors. Using a GLM to fit the predictors to the data, we estimated the response amplitudes for each condition (betas) and residual variance of each voxel’s time course. We used beta values and residual variance from the GLM to generate contrast maps comparing responses in different conditions.

*Definition of functional regions of interest (fROIs).* Spatially contiguous clusters of place-selective voxels that responded more strongly to places (scenes of houses and corridors) than to all other stimuli and were located in the collateral sulcus were defined as place-selective CoS-places (also referred to as parahippocampal place area (PPA) (Epstein and Kanwisher 1998)).

### Data acquisition in macaques (*Anatomical scans)*

*For* macaque anatomical scans, whole-brain structural volumes were acquired in a 3 T Skyra scanner while the animals were anesthetized with a combination of Ketamine (4mg/kg) and Dexdomitor (0.02mg/kg) and using a 15-channel transmit / receive knee coil. Monkeys were scanned using a magnetization-prepared rapid gradient echo (MPRAGE) sequence; 0.5 x 0.5 x 0.5 resolution; FOV = 128 mm; 256 x 256 matrix; TR = 2700 ms; TE = 3.35 ms; TI = 859 ms; flip angle = 9°). 3 whole-brain T1-weighted anatomical images were collected in each animal.

*Reconstruction of cortical surfaces.* Each animal’s T_1_ images were also co-registered and averaged. Each monkey’s average anatomical volume underwent semi-automated cortical surface reconstruction using FreeSurfer. To ensure high accuracy, skull stripping and white matter masks were first manually segmented by an expert then passed into FreeSurfer’s autosegmentation pipeline. If poor segmentations were detected, the white matter mask and control points were edited and surface reconstruction was rerun until corrected. To fix segmentation errors, average anatomical volumes were manually edited to improve the grey/white matter contrasts and to remove surrounding non-brain structures (e.g., sinuses, arachnoid, and dura matter). For all monkeys, autosegmentation of our main region of interest (the occipito-temporal sulcus) was determined to not require manual editing of the anatomical volumes.

*Functional scans.* During functional scans, monkeys were alert, and their heads were immobilized using a foam-padded helmet with a chinstrap. Monkeys were rewarded with juice for maintaining a central fixation within a 2° window. Gaze direction was monitored using an infrared eye tracker (ISCAN). Monkeys were scanned in a 3 T TimTrio scanner with an AC88 gradient insert using 4-channel surface coils (custom made by Azma Mareyam at the Martinos Imaging Center). We used a T2*-sensitive gradient echo planar pulse sequence with a repetition time (TR) of 2s, echo time (TE) of 13 ms, flip angle of 72°, slice acceleration (ipat) = 2, 1 mm isotropic voxels, matrix size 96 x 96 mm, 67 contiguous sagittal slices. To enhance contrast (Leite et al. 2002; Vanduffel et al. 2001), we injected 12 mg/kg mono-crystalline iron oxide nanoparticles (Feraheme, AMAG Pharmaceuticals, Cambridge, MA) in the saphenous vein just before scanning. Monkeys participated in 18-49 runs of a functional localizer experiment with 20s stimulus blocks with 20s of a neutral gray screen between image blocks. Monkeys viewed gray-scale stimuli, which were blocked by category. Images consisted of four categories: scenes (familiar laboratory scenes), faces (mosaics of monkey faces on a pink-noise background), bodies (mosaics of headless monkey bodies on a pink-noise background), and inanimate objects (mosaics of objects on a pink-noise background). Each image subtended 20° x 20° of visual angle and was presented for 0.5s. Each category block was presented once per scan and were presented in counter-balanced order. All images were equated for spatial frequency and luminance using the SHINE toolbox (Willenbockel et al. 2010).

*Analysis of functional data.* We analyzed data from the functional localizer experiment as previously reported (Arcaro and Livingstone 2017) using Analysis of Functional NeuroImages (AFNI (Cox 1996)), SUMA (Saad and Reynolds 2012), JIP Analysis Toolkit (written by Joseph Mandeville), and MATLAB. For each monkey, all images from each scan session were corrected for within-run and between-run motion. Data were detrended and spatially filtered using a Gaussian filter of 2-mm full-width at half-maximum to increase the signal-to-noise ratio while preserving spatial specificity. Functional volumes were registered to high-resolution (0.5 mm) T1-weighted anatomical images using a two-step linear, then nonlinear approach (JIP Analysis Toolkit) in a two-step process. First, a 12-parameter affine registration was performed between the mean EPI image for a given session and a high-resolution anatomical image. Next, a nonlinear, diffeomorphic registration was performed. Functional data were registered to both participant-specific anatomical images collected in a separate scan session and a group average template (NIH Macaque template (NMT) (Seidlitz et al. 2018); https://afni.nimh.nih.gov/NMT). For each monkey’s data, a multiple regression analysis (AFNI’s 3dDeconvolve (Cox 1996)) in the framework of a general linear model (Friston et al. 1995) was performed. Each stimulus condition was modeled with a MION-based hemodynamic response function (Leite et al. 2002). Additional regressors that accounted for variance due to baseline shifts between time series, linear drifts, and head motion parameter estimates were also included in the regression model. Due to the time course normalization, beta coefficients were scaled to reflect percent signal change. Because MION inverts the signal, the sign of the beta coefficients were inverted to follow normal fMRI conventions of increased activity represented by positive values.

*Definition of fROIs (LPP,* lateral place patch). Functionally selective voxels were defined as clusters of voxels (> 10 adjacent voxels) that responded more to images of scenes than to images of other categories (*p* < 0.0001 FDR-corrected). A contrast to faces was not used because 3 monkeys were raised without visually experiencing faces for the first year of life. For the remaining 6 monkeys that experienced faces during early development, a contrast between responses to scenes and to faces yielded qualitatively similar results.

### Generation of sulcal depth maps

For each participant (human and macaque), FreeSurfer’s automated algorithm was used to obtain sulcal depth maps (see **Fig. 2a** for sulcal depth maps in a sample macaque and human). Sulcal depth (in mm) is measured as the distance between the inflated surface and pial surface (Fischl et al. 2000) at each vertex. It is evaluated using a signed dot product of the displacement/movement vector with the outwards pointing surface normal of the white matter surface during inflation. Thus, for a sulcus, the movement vector points outwards (towards the pial surface) and therefore, the product is positive. Comparatively, for a gyral crown, the movement vector points inwards and therefore, is negative. Each participant’s sulcal depth maps (.sulc files) from the FreeSurfer’s autosegmentation algorithm were used to obtain depth measures from anatomical parcellations of the collateral sulcus (CoS) in humans and the occipito-temporal sulcus (OTS) in macaques. We then a) evaluated the deepest point in each sulcus and b) compared the sulcal pit in the place-selective region by conducting a 2-way analysis of variance (ANOVAs) with species (human/macaque), age of subjects (child/adult), and hemisphere (left/right) as factors.

**Figure 1.**
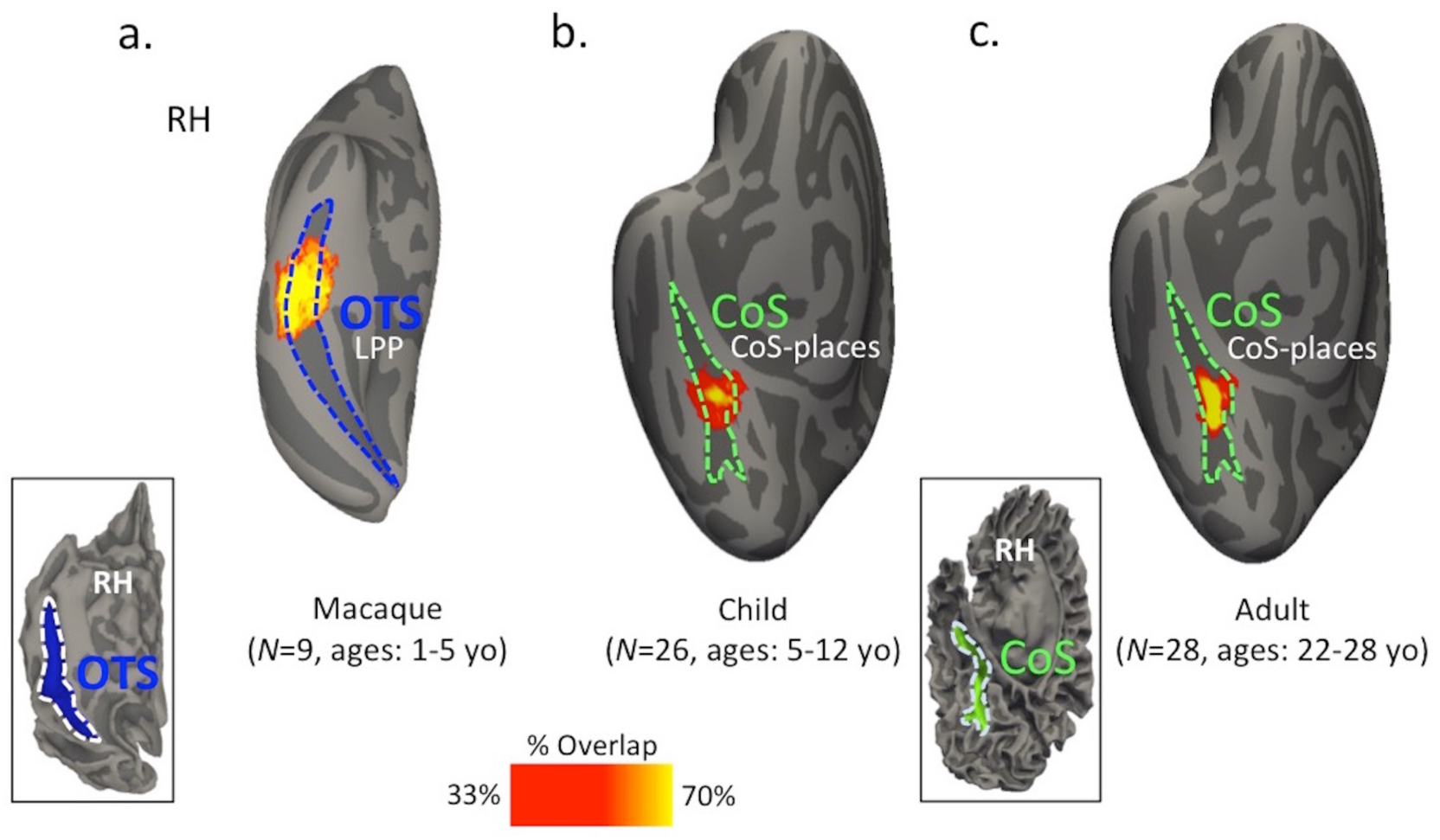
Probabilistic maps of a functionally defined place-selective region in macaques and humans are located within a sulcus in medial ventral temporal cortex (VTC) across species. (a) Inflated cortical surface reconstruction of the right hemisphere of a macaque (NIH Macaque template (NMT; Seidlitz et al. 2018; see inset for pial version). The map indicates the most probable location of the lateral place patch (LPP) in 9 macaques. The most probable location (yellow) is within the occipito-temporal sulcus (OTS; blue). (b) Inflated cortical surface reconstruction of the right hemisphere of a human anatomical template (the FreeSurfer average brain from 39 independent adult brains (Fischl et al. 1999)) showing the probabilistic location of CoS-places (also known as the PPA; Epstein and Kanwisher, 1998) in 26 children. The most probable location (yellow) of CoS-places is within the collateral sulcus (CoS; green). (c) Same as (b), but for 28 human adults. Like children, the most probable location (yellow) of CoS-places is within the CoS (green). Warmer colors in the probabilistic maps represent stronger overlap across participants within a group. Abbreviations: *RH:* right hemisphere*, CoS*: Collateral sulcu*s, OTS:* Occipital temporal sulcus. Brain sizes not to scale.

**Figure 2.**
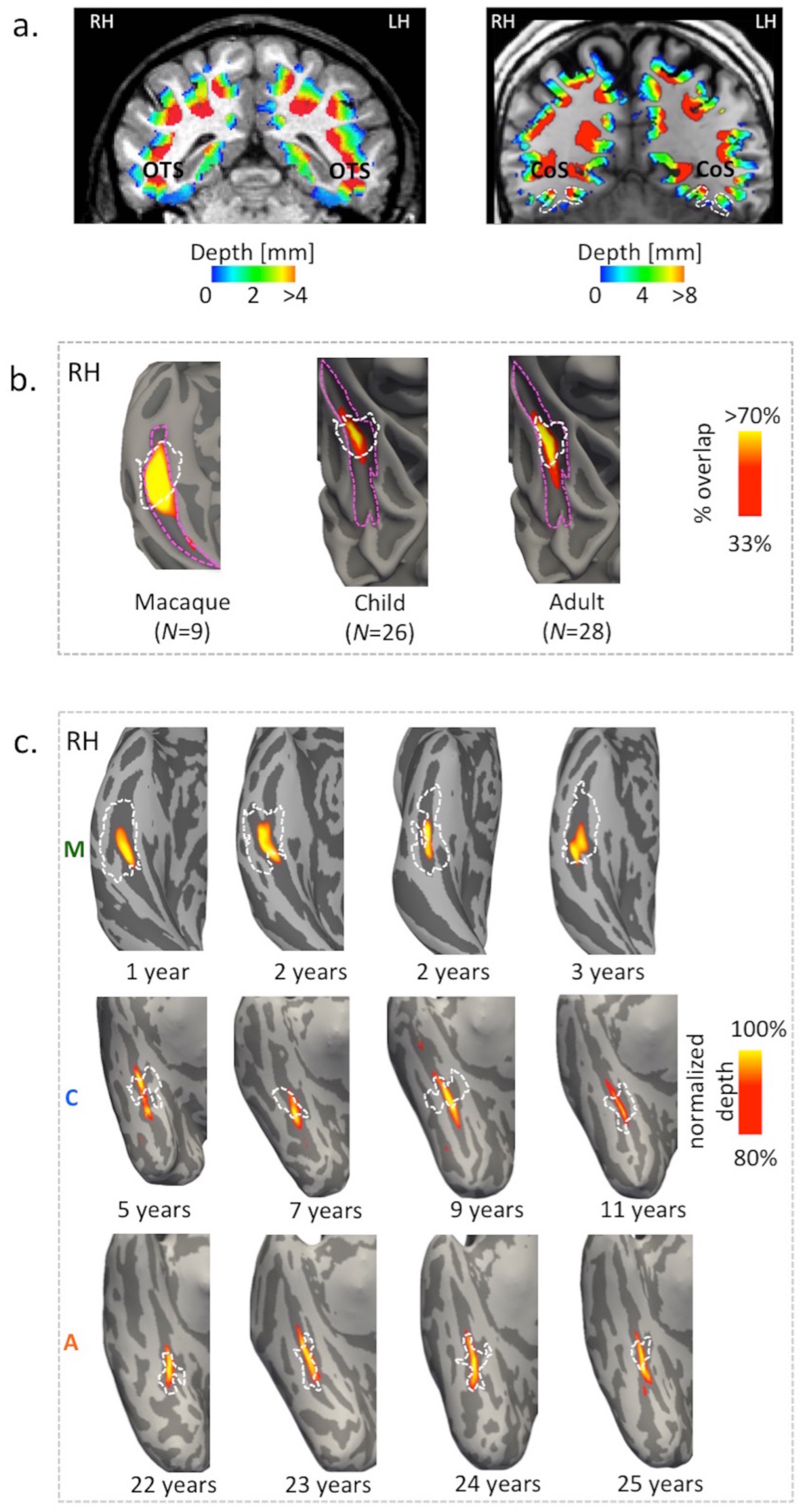
A place-selective region in medial VTC overlaps the deepest sulcal points in macaques, children, and adults. (a) Coronal slices showing depth maps obtained from FreeSurfer’s autosegmentation algorithm in sample macaque and human adult brains (warmer colors represent deeper sulci). For simplicity, the map has been thresholded to only show positive depth values representing sulcal folds, walls, and pits. (b) Inflated cortical surfaces of a right hemisphere showing probabilistic maps (percent overlap) of sulcal depth across participants within each group (macaque, child, and adult) at each point of the occipito-temporal sulcus (OTS) in macaques and the collateral sulcus (CoS) in humans (magenta dotted outlines). Across each group, the location of the deeper points (warmer colors) of a sulcus (including the sulcal pit) overlap more than the posterior and anterior edges of the sulcus. Additionally, probabilistic locations of the place-selective region (dotted white lines obtained from maps in **Fig. 1**) overlapped with the probabilistic sulcal depth maps (warm colors) in macaques, children, and adults. (c) Four sample macaque (top row), child (middle row), and adult (bottom row) ventral temporal surfaces, showing the overlap between the functionally defined place-selective region (white dotted lines) and sulcal depth maps (warm colors). To visualize the deep points of the sulcus, we thresholded the depth maps in individual subjects at 80% depth (100% is the deepest point within the sulcus). Warmer colors indicate deeper sulcal points. All individual right hemisphere surfaces are shown in **Supplementary Figs. S1, S2,** and **S3**. Brain sizes are not to scale.

### Generation of surface area and curvature values

FreeSurfer’s segmentation algorithm was also used to generate the overall curvature of the CoS and its surface area at both pial and white matter surfaces. The overall curvature quantifies the curvature (1/radius (units)) average magnitude of the entire sulcal or gyral fold.

### Probabilistic group maps of place-selectivity and sulcal depth

To visualize the location of place-selective functional ROIs (fROIs) on the cortical sheet, we generated probabilistic group maps of CoS-places in children (N=26) and adults (N=28) separately using cortex-based alignment to the FreeSurfer average brain from 39 independent adult brains (Fischl et al. 1999) (**Fig. 1**). We then calculated the percentage of subjects in which their CoS-places fROIs overlapped at each vertex on the surface. Larger overlap across participants within each age group was indicated by warmer colors on the probabilistic maps (**Fig. 1b-c**). To visualize how the sulcal depth of the CoS varied on the cortical sheet within participants of an age group, we generated probabilistic group maps of sulcal depth of the CoS in children and adults, separately using cortex-based alignment as in the prior analysis. Here, we i) transformed each participant’s sulcal depth maps into the FreeSurfer’s average brain space, ii) normalized the transformed maps based on the deepest points of the CoS, iii) added the normalized values at each point of the surface, and iv) divided by the total number of participants in each group. Then we obtained the percentage of subjects included in each vertex on the surface. In macaques (N=9), the probability maps of place-selective fROIs (**Fig. 1a**) and sulcal depth maps of the OTS were generated using the anatomical template in macaques (NMT template from 31 independent macaque brains (Seidlitz et al. 2018)) and a procedure identical to that used for humans.

### Generation of profiles of sulcal depth and selectivity in humans and macaques

For each human participant, we generated depth and selectivity profiles along the posterior-anterior axis of the CoS. This was accomplished by dividing the CoS into N bins based on the number of the z-coordinates (representing the posterior-anterior axis) along the length of the CoS. For each z-coordinate, we averaged the depth/selectivity measures along the x- (lateral-medial) and y- (superior-inferior) planes to obtain depth and selectivity profiles along the CoS. Specifically, we averaged all the cortical points in a plane orthogonal to the z axis for each z point. We then evaluated the structural-functional relationship by correlating depth and selectivity profiles in each individual participant using Pearson’s correlation coefficient (R). Similar profile analyses were conducted in the macaque OTS along its posterior-anterior axis, by first converting individual selectivity and depth maps into a common macaque coordinate space. Finally, we compared the degree of correspondence across species and human development by z-transforming the correlation coefficient measures and conducting 2-way ANOVAs with factors of species, age of subject, and hemisphere.

## Results

### A functionally defined place-selective region in medial VTC is located within a sulcus across species

To examine the consistency of the anatomical location of the functionally defined place-selective region across individuals and species, we generated group probabilistic maps of place selectivity by aligning cortical surfaces from each individual brain to their respective surface templates (Materials and Methods). Results revealed that in macaques, children, and adults, the place-selective ventral visual region is consistently located within a sulcus in medial VTC. Due to the cortical expansion across evolution (Hill et al. 2010; Chaplin et al. 2013), the sulci in which the place-selective region is located differs between species. Specifically, in humans, the place-selective region, CoS-places, is located within the collateral sulcus (CoS; **Fig. 1b-c**), while in macaques, the lateral place patch (LPP) is located within the occipito-temporal sulcus (OTS, **Fig. 1a**).

### The deepest sulcal points (sulcal pit) in medial ventral temporal cortex are in similar positions across individuals

Given the consistency in the location of the place-selective region within a sulcal fold of the medial VTC, next we sought to test if the deepest points of the sulcal fold (known as the sulcal pit) are also anatomically consistent. To do so, we aligned each individual participant’s sulcal depth map to the average surface template of their respective species and generated a probabilistic map of sulcal depth across participants. This approach revealed that within each group, the locations of the sulcal pits were well aligned across participants: the maximum percentage overlap of the sulcal pit was 100% in the left and right hemispheres of macaques, 90.09% and 91.34% in the left and right hemispheres of children, and 95.08% and 94.69% in the left and right hemispheres of adults (**Fig. 2b**), respectively.

### There is a consistent anatomical-functional relationship between sulcal depth and the location of the place-selective region in humans and macaques

In both species, the sulcus, within which the place-selective region is located, extends nearly from the occipital to the temporal pole. Given the functional consistency of the place-selective regions across species (**Fig. 1**), as well as the anatomical consistency of the deepest sulcal points across individuals within each species, we examined the overlap between each place-selective region and sulcal depth map across groups, as well as in each individual.

In all groups (macaques, children, and adults) the place-selective region overlaps the deepest points of the sulcus (**Fig. 2b**, white dotted line on probabilistic sulcal depth maps). Importantly, this structural-functional relationship was present at the individual subject level (**Fig. 2c** for sample individual subjects, and **Supplementary Figs. S1, S2** and **S3** for all subjects). Specifically, across both left and right hemispheres in 88.8% of monkeys and in 76.85% of humans (children and adults), the sulcal pit is located near the center of the place-selective region. Interestingly, however, the structural-functional overlap was more consistently observed in human adults (85.7%) than in children (68.01%). Thus, the sulcal pit is an anatomical marker that is a good predictor for the location of place-selectivity along the anterior-posterior axis of the OTS in macaques and CoS in humans even as the degree of coupling increases from childhood to adulthood.

### Sulcal-functional relationship strengthens from childhood to adulthood, but is preserved on evolutionary timescales

To further explore the relationship between sulcal depth and place-selectivity across development and species, we generated depth and place-selectivity profiles along a posterior-anterior axis of their respective sulcus (CoS for humans and OTS for macaques, see Materials and Methods). Qualitative examination of the average profiles revealed three main findings. First, we observed that in both macaques and humans, sulcal depth (**Fig. 3a,c**) and place-selectivity (**Fig. 3b,d**) have a relative smooth profile, with a single peak (represented by an *). Second, average selectivity and average depth profiles are well aligned in macaques, children, and adults. Specifically, mean place-selectivity increases as the sulcus deepens (**Fig. 3** and **Supplementary Fig. S4** for right and left hemispheres, respectively). Further, in both adults and macaques the peak of selectivity (represented by * in **Figs. 3b,d** and **Supplementary Figs. S4b,d**) is within the deepest region in the sulcus (represented by gray bar in **Fig. 3a-d** and **Supplementary Figs. S4a,c**). Third, there are clear developmental differences across human children and adults: in children, but not adults, the mean peak place selectivity was outside, but immediately adjacent, to the sulcal pit (**Fig. 3d**, peak of the blue line vs. gray bar).

**Figure 3.**
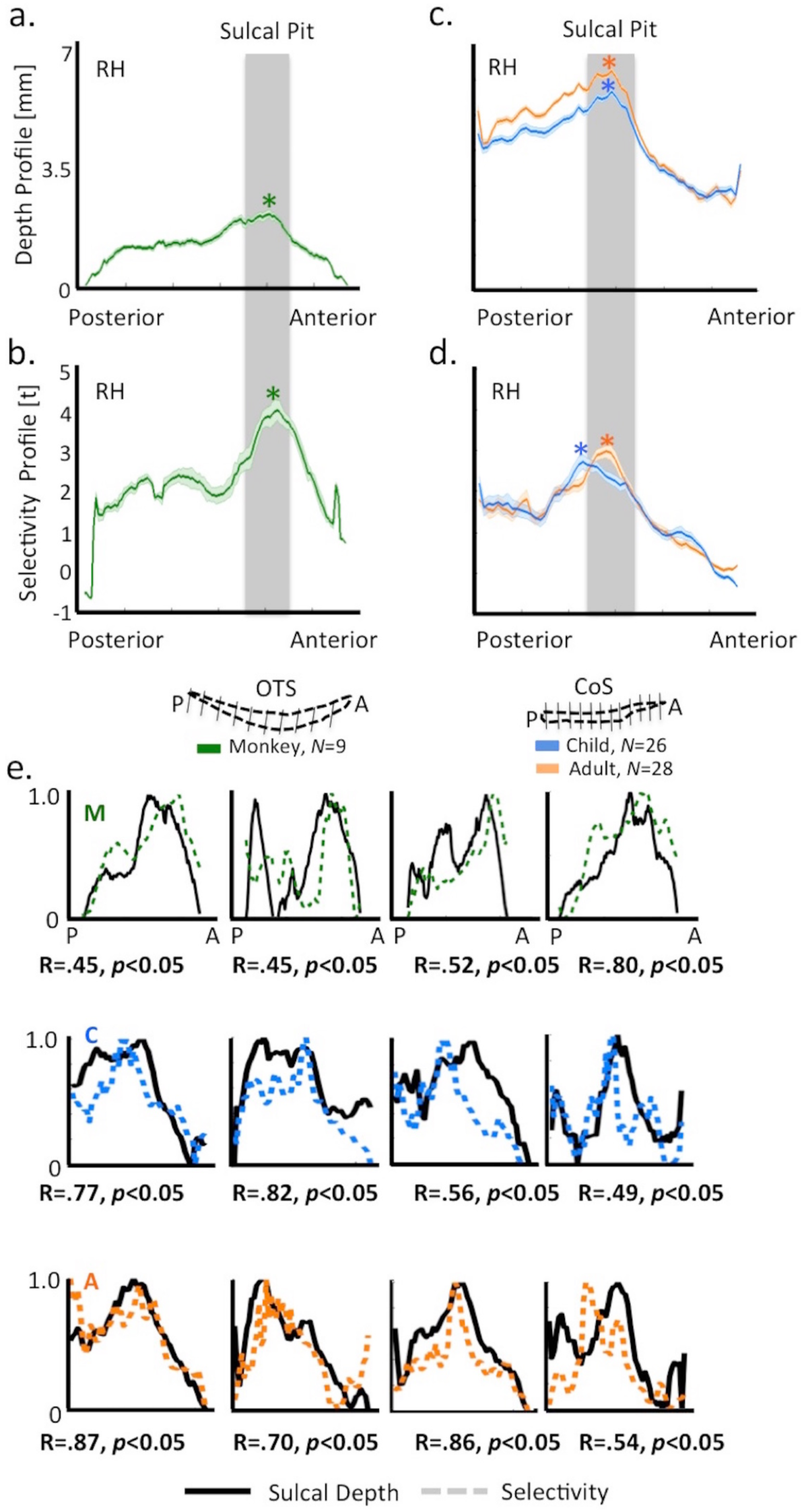
Sulcal depth and place-selectivity in medial VTC are correlated across species. (a,c) Mean (dark line) and standard deviation (lighter shading) of sulcal depth along the anterior-posterior length of the right OTS in 9 macaques (a) and the right CoS in humans (c). In (c), data are averaged across 26 children (blue) and 28 adults (orange). * represents deepest sulcal point. (b,d) Mean place-selectivity (t-value) in macaques (b) green) and humans (d). Same conventions as (a,c) Across macaques and humans, the profiles are well matched along this axis in which there is a functional peak (represented by * in b,d) around the deepest sulcal points (represented by gray bar). Left hemisphere data in **Fig. S4.** e.) Normalized depth (black lines) and selectivity (colored dotted lines) values as a function of distance along a posterior to anterior axis in four sample macaques (top row), human children (middle row), and human adults (bottom row). Correlation (R) and significance values below each subplot relate the depth and selectivity curves to one another. All individual profiles and their respective correlations are in **Supplementary Figs. S5, S6,** and **S7**.

To directly quantify this structural-functional correspondence, we calculated the correlation between each individual’s depth and selectivity profiles. This approach revealed that functional-structural profiles were significantly (R>0.28, *p*<0.05) positively correlated in 88.9% of the macaques (N=8 out of 9; R_left_: 0.4±0.12; 8 out of 9; R_right_: 0.39±0.09). In humans, the functional-structural profiles were significantly and positively correlated in 85.7% of the adults across hemispheres (N=24 out of 28; R_left_:0.49±0.05; N=24 out of 28; R_right_:0.55±0.1), as well as 61.5% of the children (N=16 out of 26; R_left_:0.35±0.06) in their left hemispheres and 73.07% of children in their right hemispheres (N=19 out of 26; R_right_:0.41±0.08; see **Supplementary Figs. S5, S6,** and **S7** for individual macaque and human functional selectivity and depth profiles, as well as their correlations).

To test if these developmental and species differences are statistically significant, we Fisher z-transformed all correlation coefficients and then statistically compared these correlations using an ANOVA. When collapsing across age groups, the correspondence between sulcal depth and place-selectivity was not significantly different between species (Macaques (M_zscore_±sem): 0.47±0.14, Humans_all_:0.55±0.05, F_1,122_=0.56, *p*>0.05; Humans_adults_:0.63±0.06, F_1,122_=2.28, *p*>0.05). Additionally, within humans, the correspondence between sulcal depth and place-selectivity significantly differed between children and adults (Children:0.43±0.08, Adults:0.63±0.06; significant main effect of age: F_1,104_=5.12, *p*<0.05). Adults showed a stronger coupling between sulcal depth and place-selectivity, as well as less variability in the structural-functional coupling compared to children. In macaques, while they did vary in age, we were unable to test for developmental effects due to the relatively small number of subjects (N=9).

Finally, to determine if the structural-functional coupling in and around the sulcal pit is driving the correlation, we used a sliding window approach to calculate the correlation of selectivity and depth into three separate bins: (1) posterior to the sulcal pit, (2) within and surrounding the sulcal pit, and (3) anterior to the sulcal pit (no sulcal point overlapped across the three bins). The correlation between sulcal depth and place-selectivity was strongest within and surrounding the sulcal pit and then decreased as the distance from the sulcal pit increased on either side of the pit for both hemispheres and species (macaque: R_mean±std_: M_close_=0.62±.3, M_away_=0.24±.5; children C_close_=0.54±.49, C_away_=0.01±.49; adults: A_close_=0.58±.45, A_away_=0.15±.48). Altogether, these results reveal that the anatomical landmark of the sulcal pit is functionally relevant within medial VTC across species. Furthermore, within humans, the structural-functional coupling between sulcal depth and place-selectivity becomes stronger and more stable from childhood to adulthood.

### What underlying mechanisms may contribute to the stronger coupling between sulcal depth and place-selectivity in medial VTC from childhood to adulthood in humans?

Prior results suggest that age-related cortical thinning in CoS-places in humans is linked to morphological changes in its curvature (Natu et al. 2019). Thus, we explored the possibility that developmental changes in sulcal fold morphology may also contribute to the structural-functional correspondence in the CoS. For example, the CoS may deepen with age and this deepening may push the functional region toward the fundus of the sulcus. Indeed, comparing CoS depth across age revealed that the CoS is deeper in adults than in children (main effect of age in a 2-way ANOVA with factors age of subject (child/adult) and hemisphere (left/right): F_1,104_=15.59, *p*<0.001, **Fig. 4a,b**). On average, the depth of the CoS sulcal pit in human adults is 10.5±0.18mm, which is ∼8.5% deeper than in human children (9.65±.23 mm, **Fig. 4b**). Intriguingly, this is not the case for all sulci. For example, in both children and adults, the deepest point is found in the insula, and there is no development in this sulcal pit from age 5 to adulthood (no main effect of age of subject: F_1,104_=3.7, *p*>0.05).

**Figure 4.**
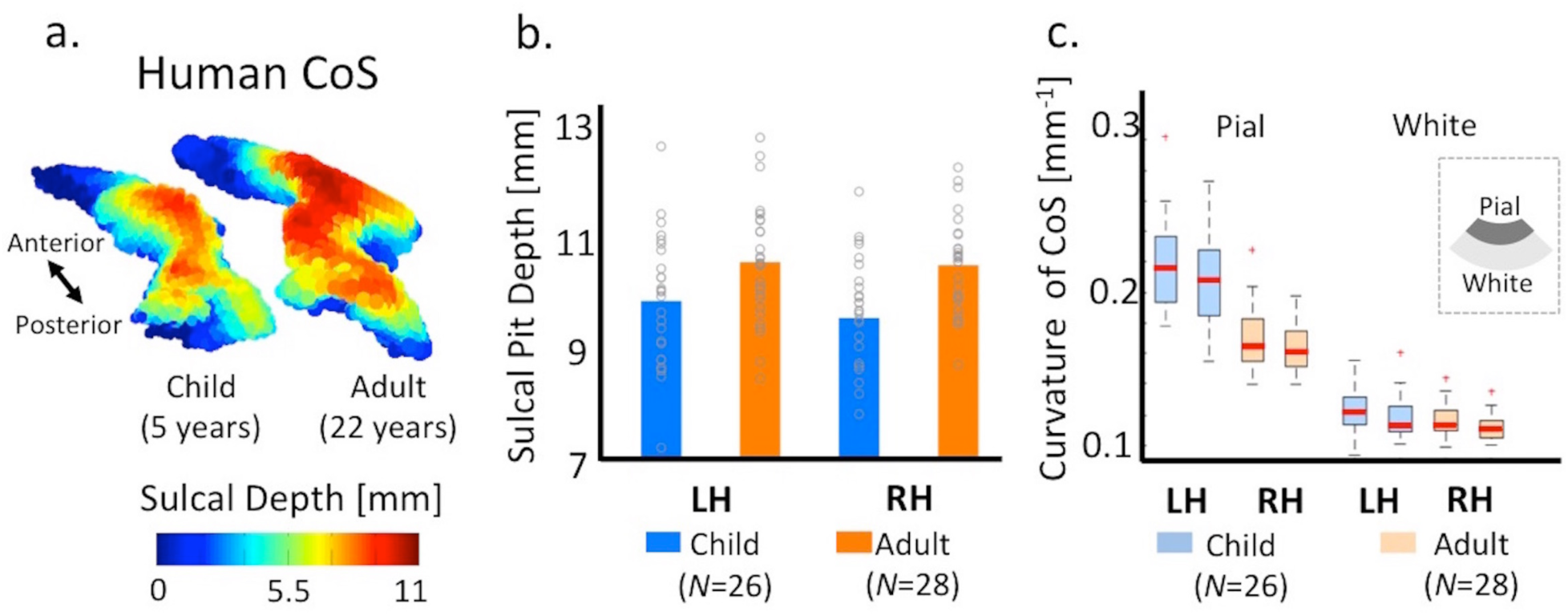
Morphological changes occur in the CoS during childhood development. (a) Three-dimensional sulcal depth maps of the collateral sulcus in an example child (5 years old), and adult (22 years old) showing differences in depth (represented by increasing height of the 3D maps). (b) Mean depth of place-selective sulcal pit in children (blue) and adults (orange). *Gray circles:* individual subjects. (c) Box plots showing median (thick red line), 25^th^, and 75^th^ percentile (box), and range (whiskers) of curvature of the CoS at the pial and white matter surfaces in children (blue) and adults (orange). *LH:* left hemisphere. *RH:* right hemisphere.

To further explore changes to the sulcal morphology that occur with this deepening, we examined if mean curvature and surface area in the upper compared to the lower cortical surfaces of the CoS also develop from childhood to adulthood. To do so, we measured the mean curvature of the CoS at the pial (upper) and white (lower) matter surfaces as prior studies suggest that differential changes in the upper versus lower cortical surfaces may result in gyrification and sulcal deepening of the cortex (Richman et al. 1975; Ronan et al. 2004). This approach revealed that the curvature at the pial surface is greater than that at the white matter surface in the CoS across age groups (F_1,208_=764.9, *p*<0.001, **Fig. 4c**) and also, that curvature develops with age. Specifically, while there is no development of the curvature at the white matter surface, the overall mean curvature at the pial surface decreased from childhood to adulthood (significant interaction between age and surface type: F_1,208_=56.03, *p*<0.001, **Fig. 4c**). Additionally, there was also significant age-related surface area expansion of both pial and white matter surfaces across hemispheres (main effect of age; F_1,208_=18.77, *p*<0.001). Together, these results demonstrate that differential morphological changes occur at the pial and white matter surfaces of the collateral sulcal fold in humans from childhood to adulthood.

## Discussion

Using comparative fMRI in macaques and humans (both children and adults), we examined the relationship between deep sulcal points and functionally-specialized regions that are critical for processing visual scenes and places in medial ventral temporal cortex (VTC). Our results revealed three main findings. First, place-selective regions in medial VTC overlap the deepest points (e.g., sulcal pits) along the OTS in macaques and the CoS in humans (**Figs. 1, 2**). Second, more broadly, there is a strong relationship between sulcal depth and place-selectivity across age groups and species in which deeper points have higher place-selectivity in the human CoS as well as in the macaque OTS; nevertheless, the strongest correlation between depth and selectivity occurred at the location of the sulcal pit and surrounding cortex in the sulcal fold (**Fig. 3**). Third, the coupling between sulcal pits and place-selective regions in medial VTC strengthens from childhood to adulthood (**Fig. 3**). Mechanistically, we propose that sulcal deepening and differential changes in the cortical curvature of the pial and white matter surfaces contribute to the strengthening of this sulcal-functional coupling from childhood to adulthood (**Fig. 4**). In the sections below, we discuss our findings in the context of (i) sulcal features as functional landmarks in high-level visual cortex in gyrencephalic brains and (ii) the role of sulcal deepening and cortical surface expansion as underlying mechanisms contributing to the observed sulcal-functional coupling in medial VTC across development and species.

### Sulcal features as functional landmarks in high-level visual cortex in gyrencephalic brains: Methodological, behavioral, and theoretical implications

Our results showing that there is a tight correspondence between the location of sulcal pits in medial VTC and place-selectivity in both macaques and humans, as well as in children and adults, is surprising because these place-selective regions are located in high-level visual cortex. Previous theories of cortical folding (Richman et al. 1975; Van Essen 1997; Welker 1990; Armstrong et al. 1995; Toro and Burnod 2005; Zilles et al. 2013) would predict a tight sulcal-functional relationship between a primary sulcus in visual cortex (i.e. the calcarine sulcus) and a primary visual area (V1). It is unclear that such sulcal-functional relationships would exist in high-level visual regions where sulcal folds are more complex and vary across individuals and primate species. Thus, these theories would not predict that a) a single point within the sulcal fundus located in what is considered one of the last stages of the visual processing hierarchy would predict the strongest selectivity of a high-level region selective for visual scenes, b) this sulcal-functional coupling would occur in both macaques and humans, or c) this sulcal-functional coupling would strengthen from childhood to adulthood - each of which we show in the present study. This latter point is particularly striking as one might expect structure-function relationships to be established early in development (potentially by genetic markers) and thus, largely fixed.

The combination of these results further illustrates the utility of sulcal features as functional landmarks in high-level visual cortex in gyrencephalic brains. For example, previous research in humans showed that a) the anterior tip of the mid-fusiform sulcus accurately identifies a region selective for faces (Weiner et al. 2010; Weiner et al. 2014), as well as b) a sulcal intersection of the collateral sulcus and the location of a region selective for visual places and scenes (Weiner et al. 2018). Here, we extend this structural-functional relationship to a specific morphological feature of sulcal depth, which is consistent with recent work showing that a sulcal pit within the human STS predicts the location of a functional region processing human voices (Bodin et al. 2018). Together, these findings suggest the possibility that a subset of functional regions in gyrencephalic brains are likely identifiable simply from sulcal depth maps instead of functional localizer experiments, which can be examined in future work.

Additional future work will likely build on this correspondence between sulcal features and functional selectivity to also incorporate aspects of behavior and cognition. For instance, sulcal interruptions in human lateral VTC have been shown to predict the location of regions selective for words as well as correlate with reading ability (Cachia et al. 2017). Furthermore, the depth of sulcal pits is correlated with different aspects of cognition such as verbal IQ (Im et al. 2011) and differences in the cortical depths and morphology of sulci are also well-known markers for developmental neurocognitive disorders such as autism (Auzias et al. 2014; Brun et al. 2016). Finally, significant hemispheric asymmetries in the frequency and distribution of sulcal pits in the superior temporal sulcus (STS) are considered to be human-specific cortical landmarks of language and socio-communication abilities (Im et al. 2010; Auzias et al. 2015; Leroy et al. 2015).

Given this growing body of work showing a relationship between sulcal depth and functional cortical representations, as well as sulcal depth and human cognition, we believe that our results are theoretically meaningful rather than epiphenomenal. Nevertheless, the commonalities between the functional arealization of visual cortex more generally across species despite vast differences in brain size, cortical folding, and gyrification (Van Essen 1997; Rosa 2005; Hill et al. 2010; Zilles et al. 2013; Arcaro and Kastner 2015) cannot be ignored as they implicate that sulci may not be necessary for the development of category-selective regions. Thus, while it is beyond the scope of the present study, future theoretical and modeling work of cortical folding relative to functional areas across species should also consider the functional role of sulcal pits in the emergence, development, and evolution of functionally-specialized areas specifically in gyrencephalic brains.

### Sulcal deepening and cortical surface expansion: Mechanistic explanations across development and species

Mechanistically, we propose that sulcal deepening and cortical surface expansion are parsimonious principles contributing to the sulcal-functional coupling observed in the present study across development and species. Developmentally, we propose that areal expansion and differential developments of the cortical curvature as the CoS deepens with age likely push the functionally active neurons closer to the sulcal fundus, thereby reducing the inter-neural distances and stabilizing the relationship between sulcal depth and place-selectivity in adulthood. Recent findings showing that place-selective cortex thins from childhood to adulthood complement this mechanistic explanation (Natu et al. 2019). Specifically, the thinning of place-selective cortex is likely linked to the morphological developments identified here in which the sulcal fold stretches and deepens leading to the shift in location of CoS-places from childhood to adulthood.

Evolutionarily, we found that the coupling between place-selectivity and sulcal depth in medial VTC was just as strong in monkeys as in humans (**Fig. 3**). This is quite surprising given that the size of the human brain is ∼15 times that of the macaque brain (**Fig. 5a**) and that other category-selective regions, such as those selective for faces, are located in different anatomical locations across species. For example, two face-selective regions are located on the fusiform gyrus (FG) in human VTC and macaques do not have an FG. Thus, the face processing system is shifted more dorsolaterally in macaques compared to humans (Weiner and Grill-Spector, 2015; Pinsk et al., 2009; Tsao et al., 2008; Arcaro and Livingstone, 2017; **Fig. 5b**). Despite this major difference between species, the topological positioning of face- and place-selective regions relative to shallow and deep sulci is consistent: two face-selective regions (ML and PL in macaque; mFus-faces and pFus-faces in human) are located adjacent to a shallow sulcus (posterior middle temporal sulcus in macaque; mid-fusiform sulcus in human), while a place-selective region (LPP in macaque; CoS-places in human) is located within a deep sulcus (OTS in macaque; CoS in human) across species (**Fig. 5b**).

**Figure 5.**
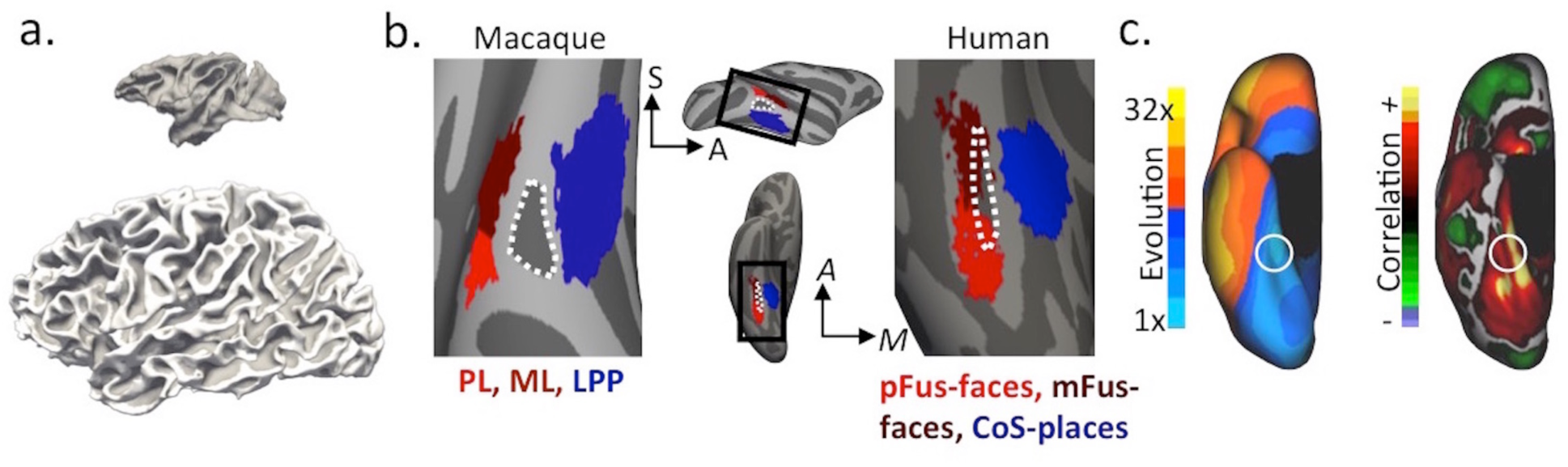
The topology of place- and face-selective regions relative to shallow and deep sulci is preserved in VTC despite significant differences in brain size and macroanatomy between macaques and humans. (a) Sagittal views of a sample macaque and a human brain (to scale) showing that human brains are ∼15 times bigger than the macaque brain. (b) Middle, Top: Lateral view of the macaque cortical surface (NMT, macaque template, (Seidlitz et al. 2018)). Black rectangle: Zoomed portion illustrated in the left image showing the macaque face-selective patches ML (maroon) and PL (red). Lateral place patch (LPP) is illustrated in blue. Each patch is the probabilistic location across 9 macaques. A shallow sulcus (posterior middle temporal sulcus, pmts; dotted white outline) divides ML and PL from LPP. Middle, Bottom: Ventral view of the human cortical surface (FreeSurfer average template from 39 adults (Fischl et al. 1999)). Black rectangle: Zoomed portion illustrated in the right image showing the human face-selective regions mFus-faces (maroon) and pFus-faces (red) relative to CoS-places (blue). Each patch is the probabilistic region across 12 adults from Weiner et al. 2017. A shallow sulcus (mid-fusiform sulcus, MFS; dotted white outline) divides mFus-faces and pFus-faces from CoS-places. (c) Left: Image of the human brain (from the PALS-B12 atlas) adapted from Hill et al. (2010). Map shows regional evolutionary cortical expansion between an adult macaque and the average human adult on the right ventral surface. White circle: Approximate location of CoS-places examined in humans in the present study. Regions in medial VTC show minimal cortical expansion compared to the remainder of cortex, while anterior-lateral VTC (warm colors) exhibits much more expansion compared to medial VTC. Right: The same cortical reconstruction of the right hemisphere with a correlation map between evolutionary and postnatal cortical expansion (image from Hill et al. 2010). Regions in medial VTC show a positive correlation across evolutionary and postnatal expansion patterns while regions in anterior-lateral VTC show little or a negative correlation. *A:* anterior*; S:* superio*r; A; M:* medial.

We propose that differences in evolutionary and postnatal cortical expansion can explain the differences and similarities in the cortical locations of face- and place-selective regions across species. Specifically, prior studies show that both postnatal and evolutionary cortical surface expansions are correlated in medial VTC (Hill et al. 2010; **Fig. 5c**). Notably, this cortical location has relatively less evolutionary expansion compared to the rest of cortex, which likely explains the coupling between sulcal depth and place-selectivity in medial VTC across species. On the contrary, anterior-lateral VTC shows no or a negative correlation between postnatal and evolutionary cortical expansion, as well as greater evolutionary expansion compared to medial VTC (**Fig. 5c**). These evolutionary anatomical differences likely contribute to the fact that category-selective regions are located on different anatomical structures across species. Furthermore, postnatal anatomical differences between lateral and medial VTC also align with the protracted development of face-selective regions in lateral VTC and the early emergence (Deen et al., 2016) and development of place-selective regions in medial VTC in humans (Golarai et al. 2007; Golarai et al. 2010; Gomez et al. 2017; Natu et al. 2019). Future work will determine if this postnatal expansion also contributes to greater individual differences in the location of face-selective regions compared to place-selective regions across age groups and species. Additionally, future work will also determine if the developmental mechanisms underlying the sulcal-functional coupling identified here in medial VTC are also comparable across species and if they extend to other functional features.

In conclusion, our study provides novel evidence of a homologous macroanatomical-functional relationship between species in which sulcal depth in medial VTC predicts place-selectivity in high-level visual cortex in both humans and macaques. This sulcal-functional relationship tightens from childhood to adulthood. Our study has broad impact as it can be used to provide deeper understanding of finer anatomical-functional relationships in other functional domains across sulcal and gyral folds, and more broadly, to examine these relationships in atypical human brain development as well as other primates.

## Conflict of Interest

The authors declare no competing financial interests.

## Acknowledgements

This work has been funded by NEI grant R01 EY02231801 and R01 EY02391501 to KGS, T-32 NEI Vision Training Fellowship to VSN

## Authors’ Contributions

VSN and KSW conceived and designed the analyses relating sulcal depth and place-selectivity across species; VSN designed the human fMRI experiment, collected human fMRI data, analyzed human and macaque data, and wrote the manuscript; MA and ML designed macaque fMRI experiment and collected macaque fMRI data. MA analyzed macaque fMRI data and wrote the manuscript; MAB and JG contributed to human fMRI data collection and analysis; KGS and KSW oversaw components of the human fMRI experiment; KSW wrote the manuscript. All co-authors have read and approved the submitted manuscript.

