## SupplementaryFigures for "Sulcal depth in medial ventral temporal cortex predicts the location of a place-selective region in macaques, children, and adults"

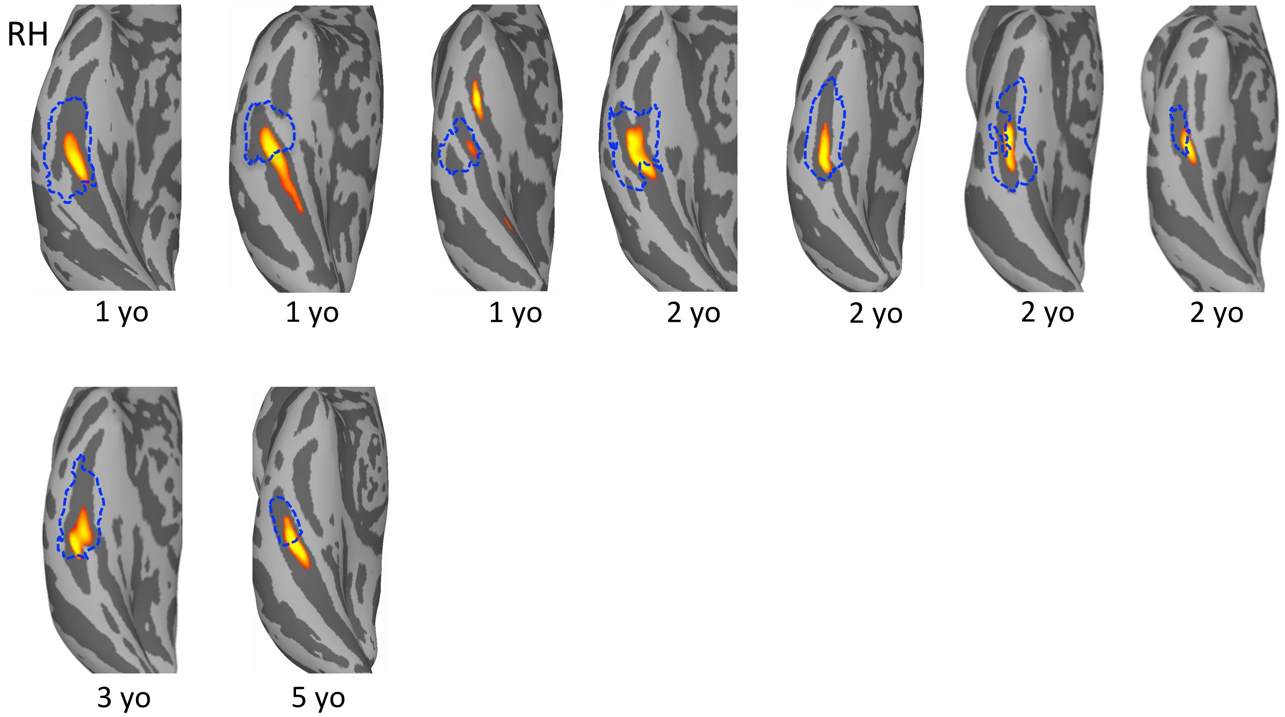


**Supplementary Figure 1**. Right ventral temporal surfaces in individual macaques, showing the overlap between the functionally defined place-selective regions (blue dotted lines) and sulcal depth maps of the occipital temporal sulcus thresholded at 80% depth of the deepest point on the sulcus. Warmer colors represent deeper areas. *Yo*: years old.

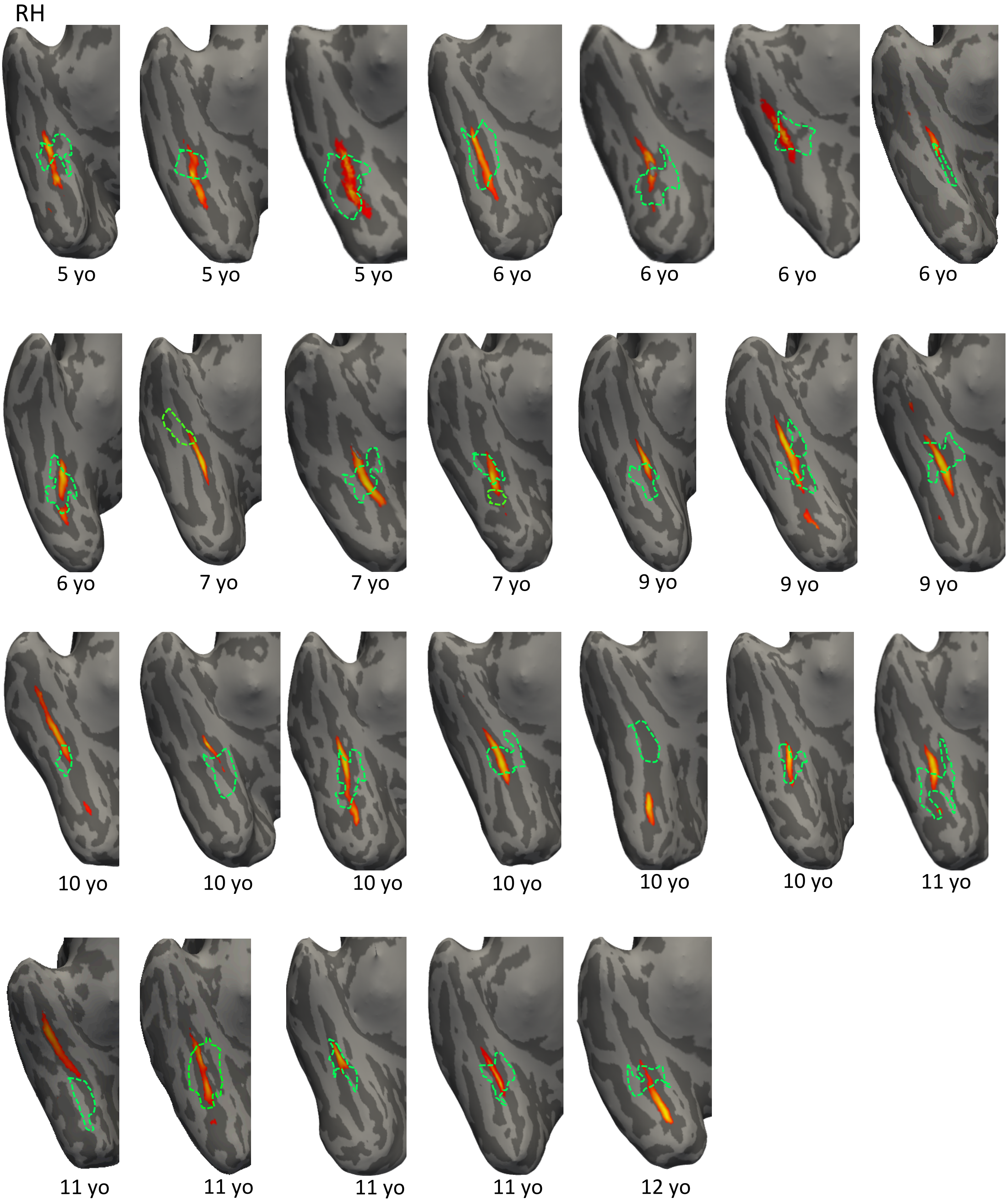


**Supplementary Figure 2**. Right ventral temporal surfaces in individual human children, showing the overlap between the functionally defined place-selective regions (green dotted lines) and sulcal depth maps of the collateral sulcus thresholded at 80% depth of the deepest point on the sulcus. Warmer colors represent deeper areas. *Yo*: years old.

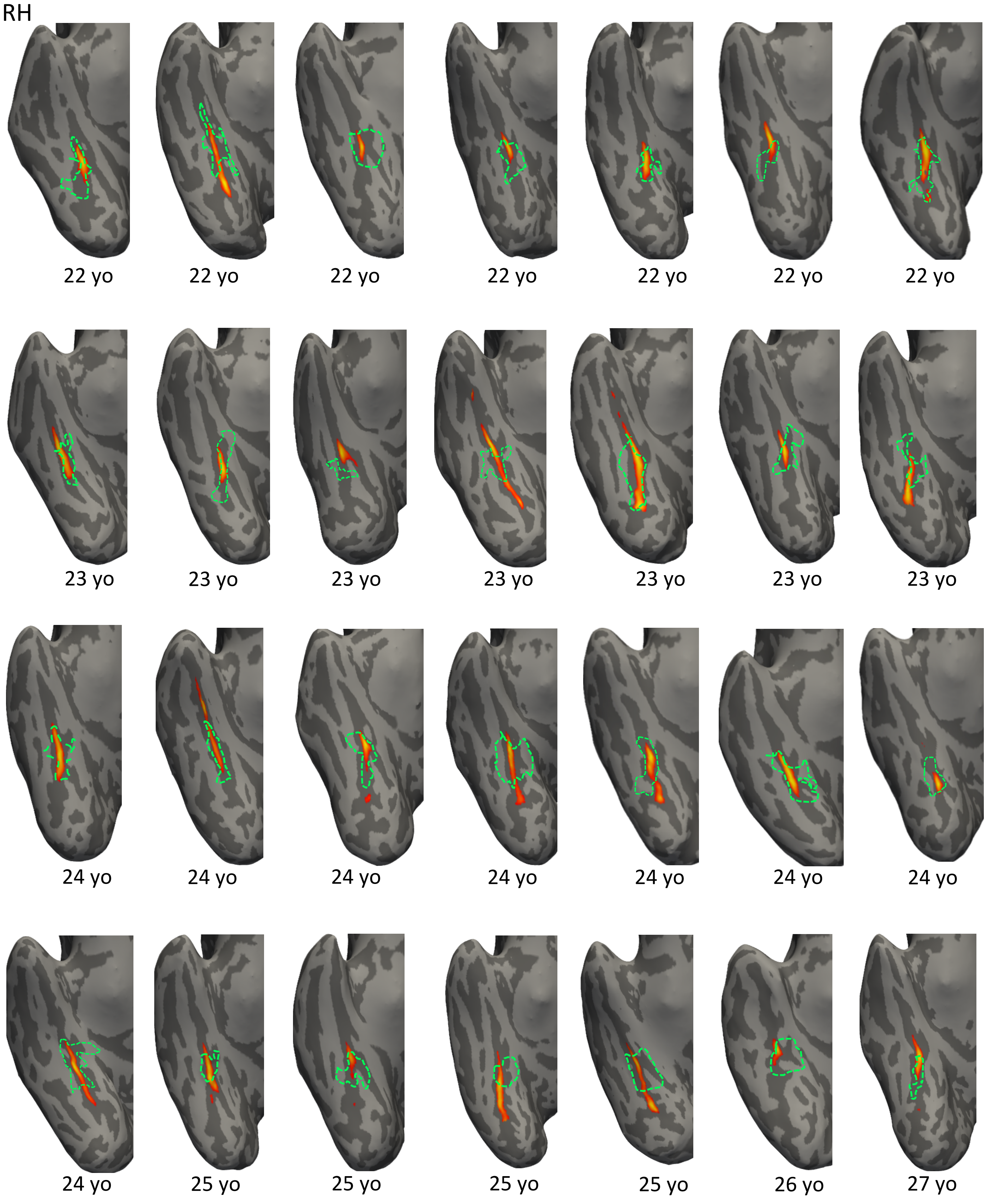


**Supplementary Figure 3**. Right ventral temporal surfaces in individual human adults, showing the overlap between the functionally defined place-selective regions (green dotted lines) and sulcal depth maps thresholded at 80% depth of the deepest point on the sulcus. Warmer colors represent deeper areas. *Yo*: years old.

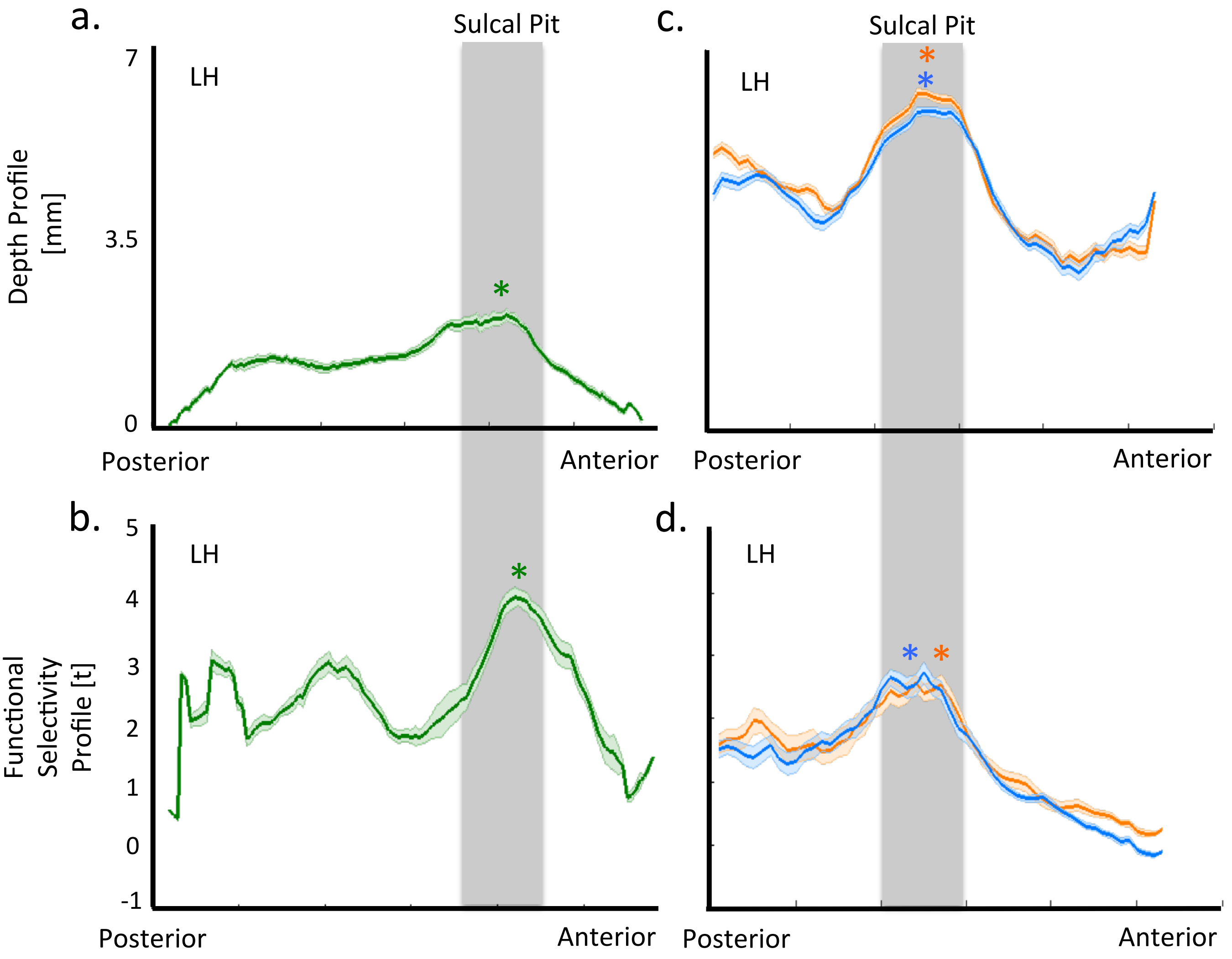


**Supplementary Figure 4. Strong correspondence between the sulcal depth and functional-selectivity profiles along the posterior-anterior axis of left place-selective sulcus in the VTC of macaques and humans.** Sulcal depth (in mm) and functional-selectivity (in t-stats) profiles of (a-b) macaques (in green), (c-d) human children (in blue) and adults (in orange) along the posterior to anterior axis of their respective place-selective sulci in VTC in the left hemisphere. Across all macaques and humans the profiles are well matched along this axis, that is functional selectivity tends to at its highest where the sulcal fold is the deepest (represented by *). (LH: Left Hemisphere).

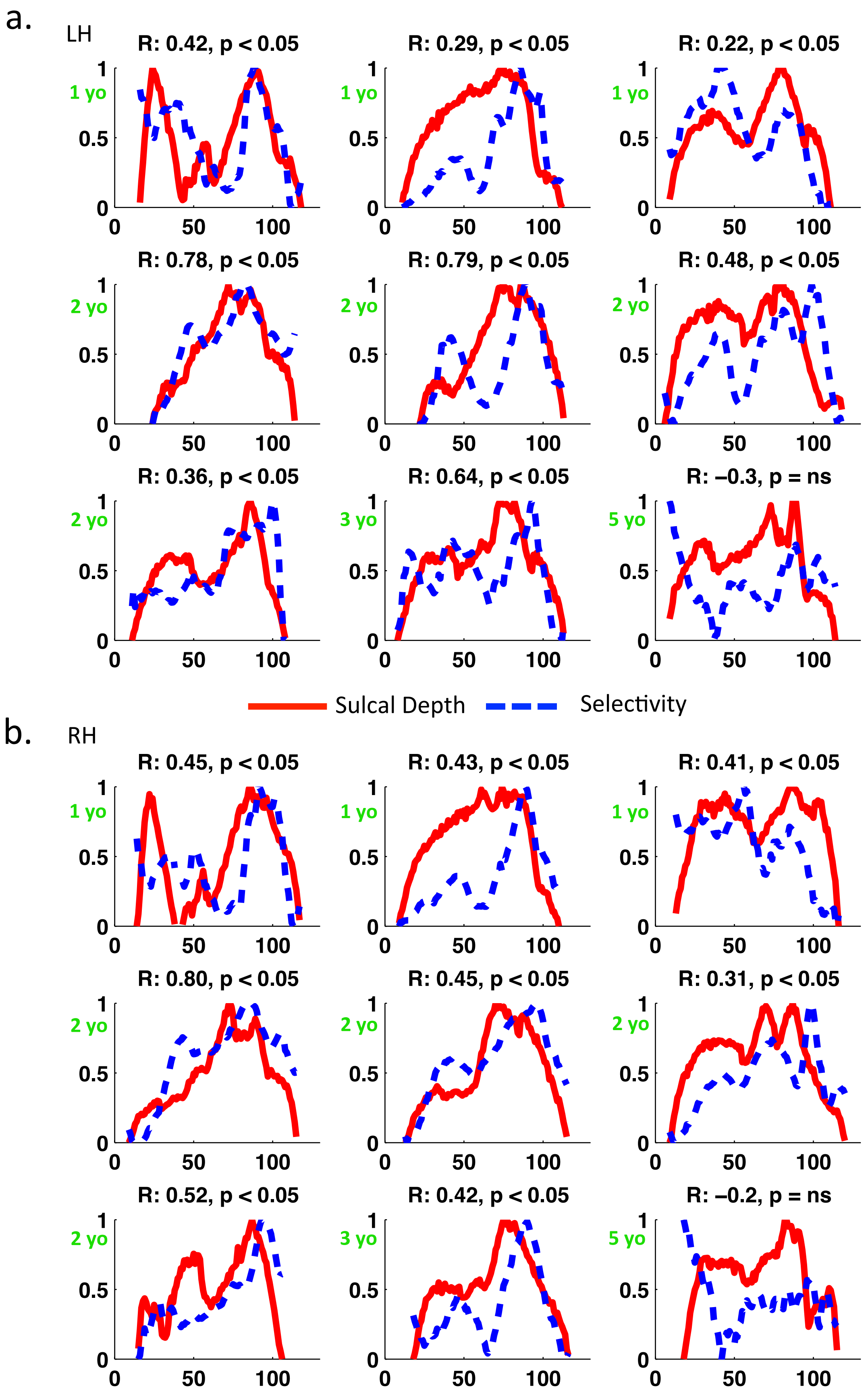


**Supplementary Figure 5. Strong relationship between the profiles of the sulcal depth and functional selectivity of place-selective sulci in macaques.** a. Figure shows the profiles of sulcal depth (red line) and place-selectivity (blue dotted line) in macaques ($N$=9) in left hemisphere. b. same as in a. in right hemisphere. Age of subject is shown in green on the left side of each graph. *LH*: left hemisphere, *RH*: right hemisphere). *Yo*: years old.

**LH**

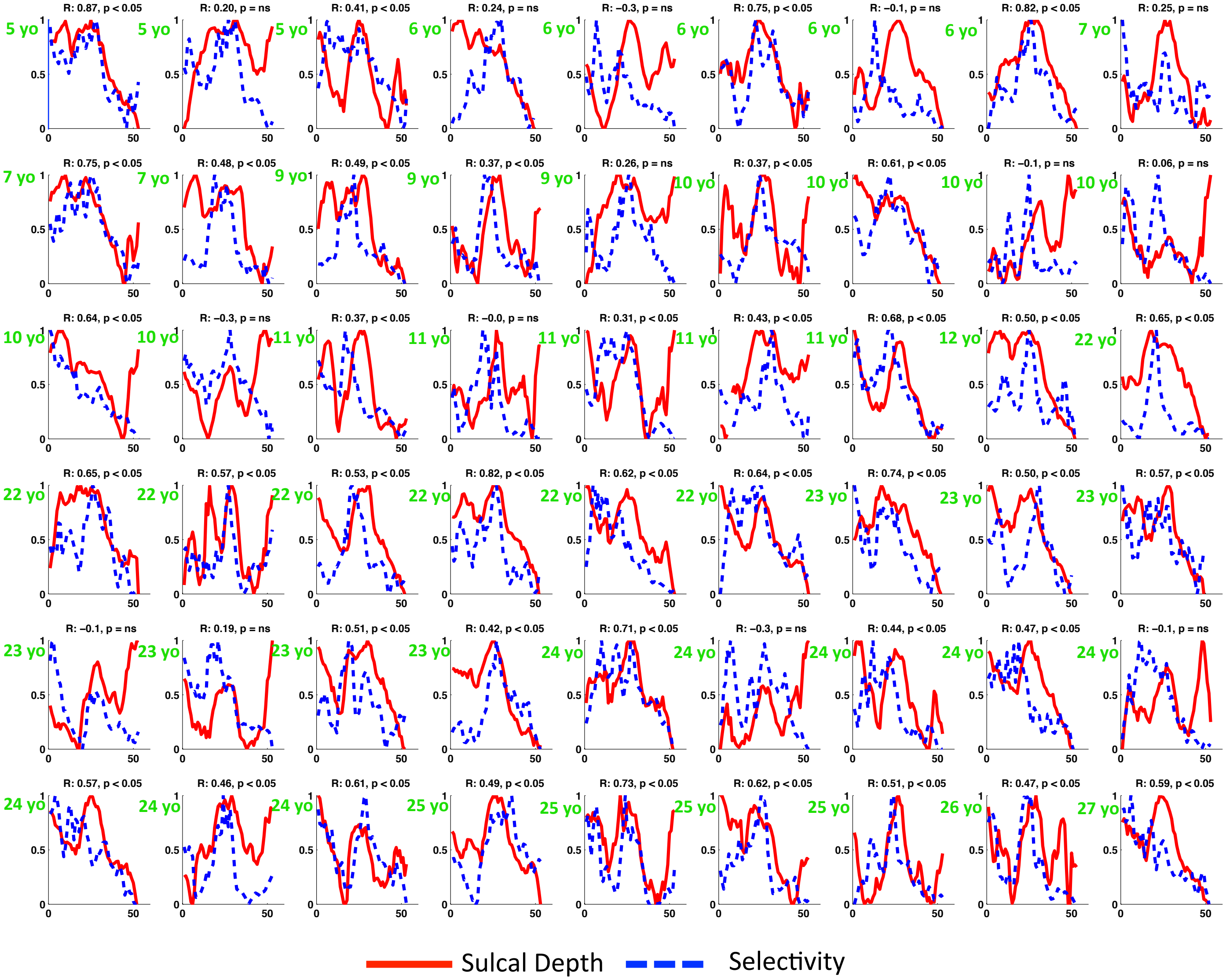


**Supplementary Figure 6. Strong relationship between the profiles of sulcal depth and functional selectivity of left place-selective sulci in humans.** Profiles of sulcal depth (red line) and place-selectivity (blue dotted line) in humans (*N*=54). Age of subject is shown in green on the left side of each graph. *LH*: left hemisphere. *Yo*: years old.

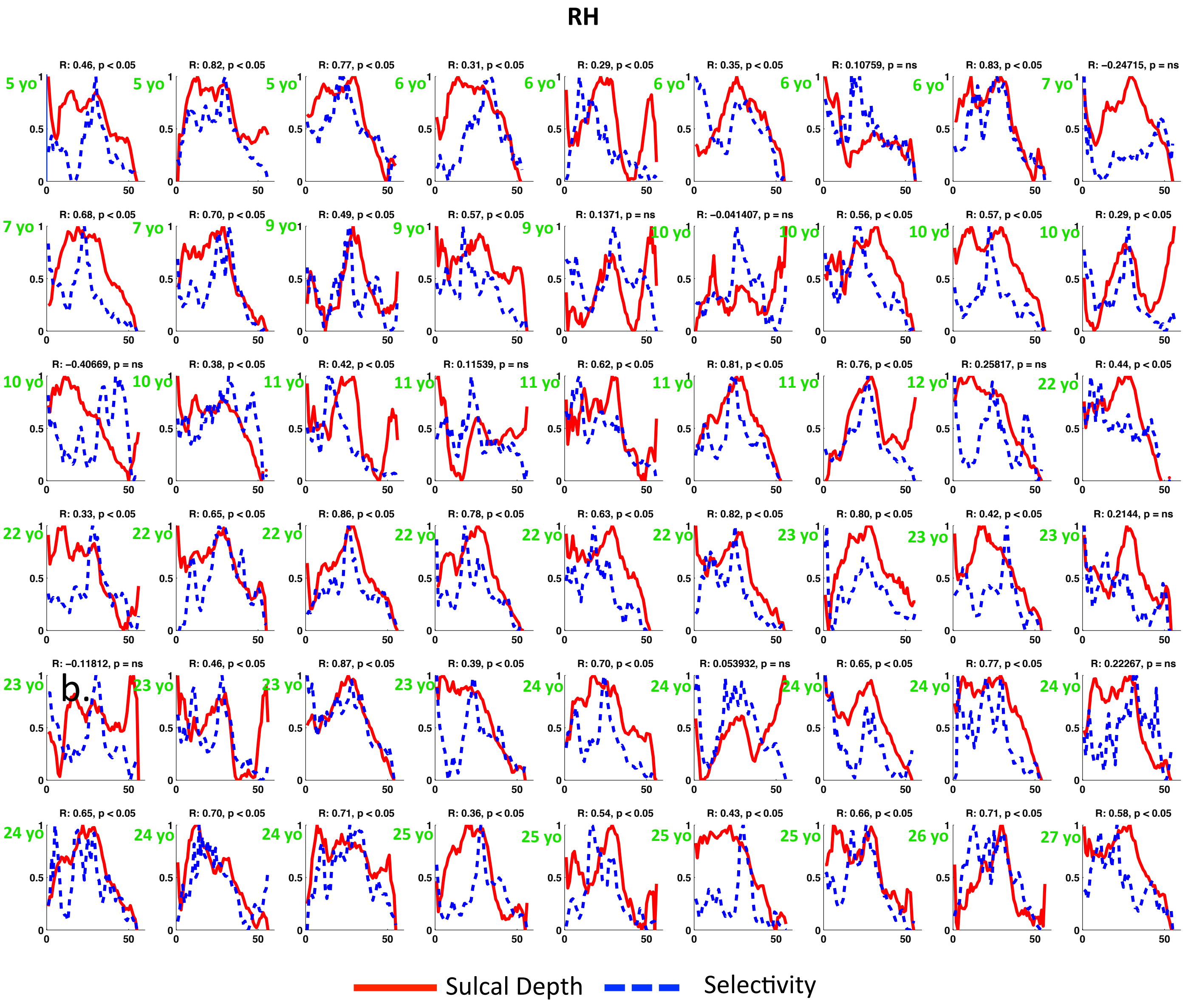


**Supplementary Figure 7. Strong relationship between the profiles of sulcal depth and functional selectivity of right place-selective sulci in humans.** Profiles of the sulcal depth (red line) and place-selectivity (blue dotted line) in humans (*N*=54). Age of subject is shown in green on the left side of each graph. *RH*: right hemisphere. *Yo*: years old.
